# A Hopfield network model of neuromodulatory arousal state

**DOI:** 10.1101/2024.09.15.613134

**Authors:** Mohammed Abdal Monium Osman, Kai Fox, Joshua Isaac Stern

**Affiliations:** Department of Neurobiology, Harvard Medical School; Kempner Institute for the Study of Natural and Artificial Intelligence, Harvard University Boston, MA, USA

## Abstract

Neural circuits display both input-driven activity that is necessary for the real-time control of behavior and internally generated activity that is necessary for memory, planning, and other cognitive processes. A key mediator between these intrinsic and evoked dynamics is arousal, an internal state variable that determines an animal’s level of engagement with its environment. It has been hypothesized that arousal state acts through neuromodulatory gain control mechanisms that suppress recurrent connectivity and amplify bottom-up input. In this paper, we instantiate this longstanding idea in a continuous Hopfield network embellished with a gain parameter that mimics arousal state by suppressing recurrent interactions between the network’s units. We show that dynamics capturing some essential effects of arousal state at the neural and cognitive levels emerge in this simple model as a single parameter—recurrent gain—is varied. Using the model’s formal connections to the Boltzmann machine and the Ising model, we offer functional interpretations of arousal state rooted in Bayesian inference and statistical physics. Finally, we liken the dynamics of neuromodulator release to an annealing schedule that facilitates adaptive behavior in ever-changing environments. In summary, we present a minimal neural network model of arousal state that exhibits rich but analytically tractable emergent behavior and reveals conceptually clarifying parallels between arousal state and seemingly unrelated phenomena.

## 1 Introduction

Neural circuits display both input-driven dynamics that are necessary for the real-time control of behavior and internally generated dynamics that are necessary for memory, planning, and other cognitive processes. These forms of activity often complement one another, as when a sensation triggers the recall of a relevant memory. However, they can also conflict, as when pleasant recollections leave a daydreamer oblivious to his surroundings. In animals, a potent regulator of this balance between intrinsic and evoked dynamics is arousal state [27, 16]. Arousal is an internal state variable that regulates the organism’s overall level of activity and sensitivity to external input. During high arousal states, weak sensory inputs can achieve system-wide control of perception and behavior. In contrast, during low arousal states, such as sleep, neural dynamics are decoupled from external input and dominated by spontaneous activity that is thought to reflect the querying of internal models [27, 21]. It has been hypothesized that arousal state exerts these effects through gain control mechanisms that suppress the recurrent connections that generate spontaneous activity and amplify the feedforward connections that convey external input, an idea consistent with the known effects of acetylcholine, a potent neuromodulator of arousal state [9, 7, 6].

In this paper, we study this interplay between arousal state, spontaneous activity, and stimulus-evoked dynamics in a continuous Hopfield network [12] embellished with a gain parameter that mimics arousal state by suppressing recurrent interactions between the network’s units [9]. We show that varying the network’s recurrent gain induces a phase transition from a low arousal state dominated by intrinsic dynamics, memory retrieval, and Bayesian priors, to a high arousal state dominated by evoked dynamics and sensory input. To build further intuition, we highlight parallels between our model’s behavior and physical phenomena studied in statistical mechanics. Turning to biology, we then discuss how the dynamics of neuromodulator release flexibly balance a tradeoff between the reliable sensory responses characteristic of high arousal states and the cognitive functions enabled by strong recurrence. In summary, we present a minimal neural network model whose analytical tractability and emergent dynamics shed light on the functional role of arousal state in neural computation.

## 2 Related Work

A rich body of prior work has explored how neuromodulatory gain control mechanisms can alter the balance between intrinsic and evoked dynamics in neural circuits. For example, [9] put forth a circuit model of olfactory associative learning in piriform cortex (very similar to the Hopfield network analyzed in the present study) in which acetylcholine determines the multiplicative gain of the circuit’s recurrent connections and explored how it interacts with synaptic plasticity. In addition, a recent study of auditory cortex has shown that modulating the gain on background inputs to excitatory cells can induce a transition from multistable to unistable attractor dynamics and explain the inverted-U relationship between arousal state and task performance in perceptual decision making [20]. Finally, in the active inference literature, previous work has argued that acetylcholine sets the balance between the likelihood and prior terms in Bayesian inference by modulating the response gain of pyramidal cells thought to convey sensory prediction errors to downstream brain areas [17].

Outside the context of arousal state, classic work on recurrent neural networks with random synaptic couplings has demonstrated that they undergo a phase transition from unistable attractor dynamics to chaos as neural gain is increased, an insight relevant to our model despite its symmetric connectivity [25]. Finally, other work has employed deterministic annealing approaches, in which neural gain is gradually increased as some computation progresses to encourage the network to settle into an attractor state representing a good solution to an optimization problem, rather than getting trapped in a local minimum, another concept that will prove to be relevant [18, 2].

## 3 Network Dynamics

In this paper, we study a continuous Hopfield network with *N* recurrently coupled units whose time varying activations are governed by the differential equation

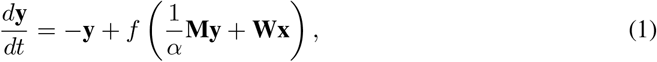

where **y** ∈ ℝ^*N*^ denotes the vector of neural activations, **x** ∈ ℝ^*D*^ denotes the stimulus pattern, **M** ∈ ℝ^*N*×*N*^ denotes the symmetric and zero-diagonal recurrent connectivity matrix, **W** ∈ ℝ^*N*×*D*^ denotes the feedforward input weights, and *f* (*·*) is an element-wise tanh nonlinearity. Due to their symmetric connectivity, networks of this form exhibit simple point attractor dynamics that are commonly used to model associative memory.

In addition to the usual variables, the dynamics are also determined by an “arousal” parameter *α* ∈ ℝ_+_ that divisively suppresses recurrent interactions between the network’s units (Figure 1). In brief, when *α* is low, recurrence is strong and intrinsic dynamics dominate, while when *α* is high, recurrence is weak and the network state is controllable by bottom-up sensory input.

**Figure 1:**
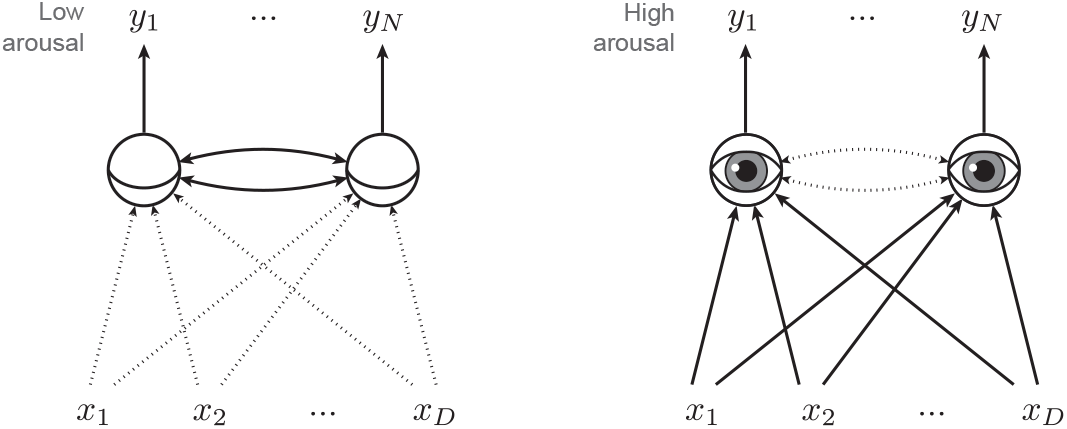
Network schematic illustrating low arousal state dominated by recurrent interactions (left) and high arousal state dominated by bottom-up sensory input (right).

More formally, as *α* approaches 0, the dynamics approach

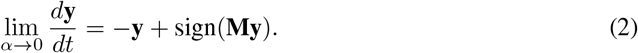

In this low arousal, sleep-like limit, sensory input has no effect on the network dynamics and the attractor states of the system correspond to the memory patterns stored in its recurrent weights. Conversely, as arousal diverges, the dynamics approach

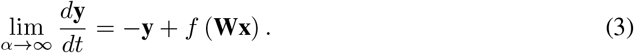

In this high arousal limit, the dynamics are completely controlled by the input, the activity state of the network exponentially relaxes towards the stimulus-evoked fixed point at *f* (**Wx**), and the system effectively reduces to a feedforward logistic function of the input.

Note that the network is multistable in the low arousal limit and unistable in the high arousal limit. As this qualitative change in the fixed point structure entails, these regimes are separated by a bifurcation (or series of bifurcations). While a closed form expression for the initial bifurcation’s critical point is unavailable when input is present, in the absence of input, it is solely determined by the largest eigenvalue of the connectivity matrix:

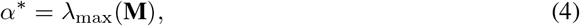

which is proven in the Appendix.

Further intuition can be obtained by considering a simple instantiation of the model consisting of two mutually inhibitory neurons (Figure 2). At a given value of *α*, to visualize the dynamics of this two dimensional system, we plot its flow field and overlay each unit’s nullcline, the set of states for which the time derivative of that unit’s activity is zero. Each nullcline is a sigmoidal curve whose steepness is determined by *α* and whose translational position is determined by the corresponding unit’s input. For a fixed input, at low *α*, the nullclines are steep and intersect at three points, resulting in two attractor states and a saddle node, while at high *α*, the nullclines are shallow and intersect at just one point, which becomes the global attractor state of the system.

**Figure 2:**
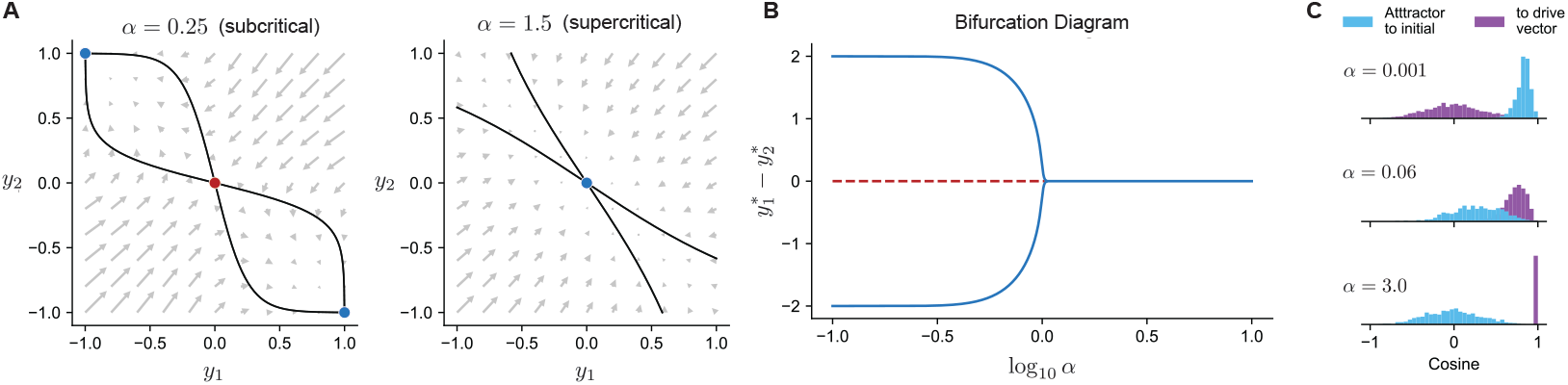
Network undergoes bifurcation from multistable associative dynamics to unistable input driven dynamics as alpha passes a critical point. (a) Phase portraits for the dynamics of 2-unit network under zero external stimulus. Black lines indicate nullclines of the dynamics, whose intersections result in the stable (blue) and unstable (red) fixed points. (b) Difference of unit activations at stable (blue, solid) and unstable (red, dashed) fixed points as a function of recurrence gain. (c) Distribution of angles between initialization point of integration and eventual attractor state for uniform random initializations and drive vectors **Wx** (purple) as well as between attractor and drive vector (blue).

## 4 Energy Function

The dynamics of the network can also be construed as descent along an energy landscape whose local minima define the attractor states. This energy landscape is defined by the Lyuapanov function

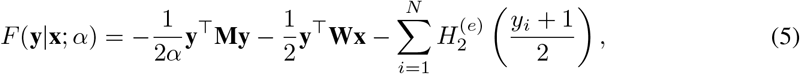

where 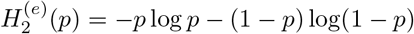 [12]. This equation for the Lyuapanov function of the network dynamics also equals the mean-field variational free energy for a probabilistic generative model called a Boltzmann machine [11, 1, 14]. While we will defer a more detailed overview of this connection to the Appendix, in brief, this equivalence allows us to cast the dynamics of the system as an approximate Bayesian inference procedure in which the (variational free) energy monotonically decreases as the system approaches an attractor state representing the locally optimal inference about the latent causes of its sensory input. In this framing, the network state **y** encodes a factorized probability distribution *q*(**z**) over the generative model’s latent variables and the energy *F* (**y** |**x**; α) quantifies (or more precisely, upper-bounds) the discrepancy between *q*(**z**) and the true Bayesian posterior *p*(**z**|**x**) [3, 5]. With these considerations in mind, we can equivalently write 5 as

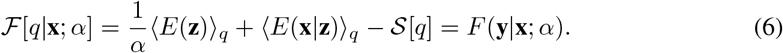

To minimize this variational free energy, the allocation of posterior probability—as represented by the activity state of the network—jointly favors states that have high prior probability (through 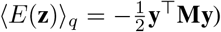 and compatibility with the data (through 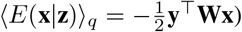, while also maximizing the entropy of the posterior (through 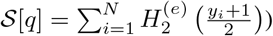. As this decomposition reveals, the arousal level *α* determines the weakness of the prior used in inference. As *α* goes to infinity, the contribution of the prior term vanishes and the posterior probability of a state merely reflects its compatibility with the data. In this limit, inference reduces to maximum likelihood estimation regularized by an entropy term, and the underlying generative model reduces to a restricted Boltzmann machine. Conversely, as *α* goes to 0, the likelihood and entropy terms become negligible and the posterior probability of a state depends only on the prior. Varying *α* therefore allows the network to interpolate between sampling from its prior and performing inference over the data.

We can gain further intuition by considering some analogies to physical systems. In the absence of input, equation 6 reduces to 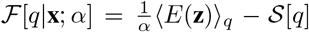and *α* functions as a temperature parameter. Correspondingly, in the high arousal regime, the system minimizes variational free energy by maximizing the entropy of its belief distribution and thereby remaining agnostic about the unobserved state of the world. In contrast, in the low arousal regime, the system’s strong priors instead favor the allocation of all belief to an arbitrary state that is likely under the generative model.

In the presence of input, temperature becomes an imperfect metaphor and *α* instead corresponds to the inverse of interaction strength in the Ising model. The Ising model is a lattice of interacting binary spin variables that represents an idealized magnetic solid experiencing the effects of temperature and an external field, and its interaction strength parameterizes the energetic cost of misalignment between neighboring spins [13, 19]. At subcritical interaction strengths, couplings within the lattice are too weak to buffer an arbitrary state against thermal fluctuations or an oppositely oriented applied field, so the solid’s magnetization passively reflects that of the world around it. Above a critical point, in contrast, strong internal interactions lead to a global symmetry breaking event in which most of the spins adopt some arbitrary orientation that is subsequently hard to reverse, even if it is misaligned with the external field. Surprisingly, this suggests that the unreliable sensory responses and internally generated imagery of low arousal states like daydreaming are analogous to the hysteresis and spontaneous symmetry breaking properties of ferromagnetic solids.

## 5 Discussion

As these results highlight, while the recurrent dynamics that produce robust pattern completion, persistent activity, and structured spontaneous fluctuations enable cognitive functions like associative recall, working memory, and imagination, they do so at the expense of reliable responses to external input. To cope with this fundamental trade-off, neuromodulatory mechanisms must therefore match the balance between intrinsic and evoked dynamics to the current needs of the organism [23, 15, 24].

On slow time scales, this balance is manifest in sleep-wake cycles, and on intermediate time scales, the alternation between daydreaming and engaged task performance has a similar flavor. On fast time scales, moment-to-moment fluctuations in levels of acetylcholine and norepinephrine—as indexed by pupil dynamics—also have analogous effects [16, 22]. For example, pupil dilation accompanies perceptual flips in the interpretation of an ambiguous stimulus like the Necker cube, potentially indicating a momentary suppression of recurrence that permits switching between attractor states [26]. Conversely, pupil constriction accompanies hippocampal and cortical replay events, suggesting low neuromodulatory tone and strong recurrence during moments of memory recall [4]. Finally, models of the theta rhythm have argued that oscillating levels of acetylcholine in the hippocampus define a fast cycle of stimulus encoding and memory retrieval [8]. Taken together, these phenomena suggest that alternation between these modes structures neural computation across scales.

We propose that the dynamics of arousal state can be understood through the lens of yet another concept from statistical mechanics: namely, annealing. Annealing is the gradual reduction of a material or search algorithm’s temperature as an optimization process unfolds, and it is premised on the idea that smoothly deforming a system’s energy landscape in this manner can enable it to relax into a globally optimal configuration rather than getting stuck in a local minimum [2]. In neural circuits such as the continuous Hopfield model presented here, neural gain essentially plays the same role as temperature in annealing algorithms [18]. Unlike more rudimentary annealers, however, animals confront ever changing environments and therefore optimization problems. As such, the brain’s neuromodulatory dynamics may constitute a remarkably rich and flexible annealing schedule that facilitates adaptive behavior and information processing under such circumstances (Figure 3).

**Figure 3:**
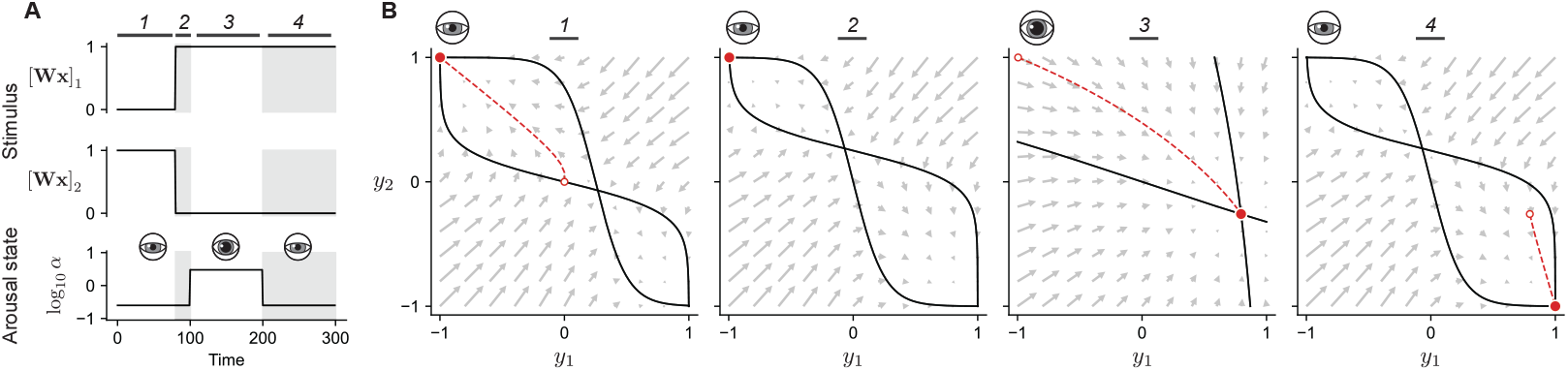
Dynamic annealing schematic: Momentary suppression of recurrence allows escape from suboptimal local minima, potentially explaining pupillary dynamics during perceptual switching. (a) Input drives to a 2-unit network and dynamic arousal state chosen to switch to a high value for a short period after stimulus switching. (b) Recurrence gain allows for dynamic control of hysteresis and annealing. Network state (red) during the example stimulus, first maintaining current activity invariant to stimulus switch (2), then adapting to new stimulus characteristics during high-arousal state (3) before collapsing to a corresponding intrinsic attractor (4).

## 6 Conclusion

In summary, we analyzed a minimal neural network model of arousal state premised on the longstanding idea that arousal state modulates cognitive and sensory processing by setting the balance between recurrent and feedforward connectivity. We showed that this model recapitulates several hallmarks of arousal state despite its simplicity and interpreted this behavior through the lenses of associative memory, Bayesian inference, and statistical mechanics. Our results shed light on the functional significance of arousal state and recurrent gain modulation in brain circuits, potentially suggesting that similar mechanisms could enhance the flexibility of neural computation in AI systems.

## Aknowledgements

Many thanks to Mark Andermann, Ila Fiete, Akshay Jaggi, Cengiz Pehlevan, Jacob Zavatone-Veth, Stelios Smirnakis, Ganna Palagina, and Gord Fishell for enlightening discussions about these ideas.

## A Deriving the Critical Point

Here we derive the critical arousal level α^*^ at which the network undergoes a phase transition (i.e. pitchfork bifurcation) from multistable to unistable attractor dynamics. In the absence of input, this occurs when the fixed point at the origin goes from being an attractor to a saddle point. To determine when this occurs, we will perform a linear stability analysis, linearizing the dynamics around the origin and determining when the largest eigenvalue of the Jacobian **J** |_**0**_ becomes greater than 0, as a function of α. The entries of the Jacobian are given by

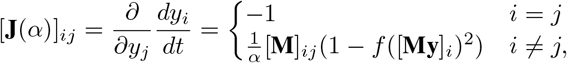

which evaluated at the origin yields

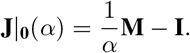

Then we see that the critical point is simply a function of this matrix’s eigenvalues:

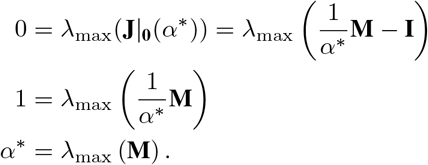

Finally, we note that since *α* is a non-negative scalar, this critical point will only lie within the relevant parameter range if, *λ*_max_(**M**) ≥ 0. As we will now show, this condition always holds because of the symmetry and zero-diagonal constraints we have imposed on **M**. First, since **M** is symmetric, all of its eigenvalues are real, and second, since the diagonal of **M** is zero,

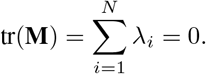

Since the eigenvalues sum to zero, either all of them equal zero or at least one is positive and one is negative. In either case *λ*_max_(**M**) ≥ 0, completing the proof.

## B Boltzmann Machines and Variational Inference

Here we review Boltzmann machines and variational inference as they pertain to the problem at hand (though see [10] and [3] for more general overviews). The Boltzmann machine [11] is a generative model consisting of *N* binary latent variables that are dubbed “causes” and collectively encoded in the “world state” **z** ∈ {*±*1}^*N*^. The couplings between these variables are summarized in the matrix 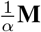, such that alignment between *z*_*i*_ and *z*_*j*_ is energetically favorable if 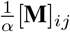 is positive and energetically unfavorable if 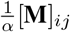. While the world state **z** cannot be accessed directly, it gives rise to an observation **x** ∈ ℝ^*D*^ via a noisy generative process parameterized by the matrix **W**. Given this observation and the parameters of the model, the goal of inference is compute *p*(**z** |**x**; *α*), the posterior distribution over world states. This probability is related to the internal energy

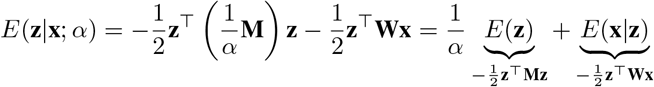

via the Boltzmann distribution

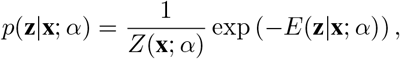

where *Z*(**x**; *α*)= Σ_**z**_ exp (−*E*(**z**|**x**; *α*)) is a normalizing constant called the partition function.

In practice, computing the true Bayesian posterior via equation (3) is infeasible because evaluating the partition function involves summing over all 2^*N*^ possible world states. This motivates the use of variational methods, a class of inference algorithms that seeks to approximate the true posterior using a family of distributions that is tractable to parameterize and optimize over [3, 5]. In this case, we turn to the mean-field approximation, which assumes that the true posterior factorizes into the product of an independent distribution for each latent variable: i.e. that

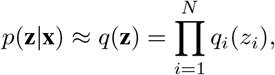

where each of the *N* factors can be parameterized by the probability it assigns a value of +1 for its associated latent, i.e. *q*_*i*_(*z*_*i*_ = +1). While the mean field approximation makes a very strong independence assumption that does not hold, one can nonetheless optimize over this family of distributions (i.e. through gradient descent) to obtain a reasonable approximation of the true posterior. This optimization amounts to minimizing a quantity called the variational free energy, which is equivalent to minimizing an upper bound on the KL divergence from the approximating distribution to the true Bayesian posterior [14, 3]. In this case, the mean field variational free energy is given by

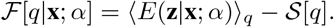

Since the variables are binary and independent, the expected world state under a candidate approximating distribution **y** = ⟨**z** ⟩_*q*_ uniquely specifies it. We can therefore equivalently write equation (4) as a function of **y** and simplify to recover the Lyapunov function of the continuous Hopfield network.

## C Numerical Simulations

To generate the plots in figures 2 and 3, we used the system’s governing equation to determine the flow fields and nullclines and simulated its dynamics using Euler’s method, a basic and widely used iterative, discrete time update rule of the form

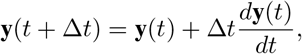

where the initial condition **y**(0) is specified by the user and the time derivative 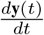 depends on the current state, arousal level, external input, and network parameters, as described in the main text 1. Dynamics for phase diagrams were calculated using two-unit networks with inhibitory connections of strength −1 between the units. Nullclines were computed analytically.

To generate angle distributions from initializations and stimuli to trajectories’ limiting behavior, we created a 10-unit network with **W** = **I** and **M** = **I − 11**^T^ to generate a densely connected inhibitory recurrent structure. Pairs (*n* = 1000) of vectors **x, y**_0_ were generated with elements uniformly distributed on [−1, 1] and stimulated dynamics were run until convergence for each arousal state.

